# Genomic epidemiology of *Mycobacterium bovis* infection in sympatric badger and cattle populations in Northern Ireland

**DOI:** 10.1101/2021.03.12.435101

**Authors:** Assel Akhmetova, Jimena Guerrero, Paul McAdam, Liliana C.M. Salvador, Joseph Crispell, John Lavery, Eleanor Presho, Rowland R. Kao, Roman Biek, Fraser Menzies, Nigel Trimble, Roland Harwood, P. Theo Pepler, Katarina Oravcova, Jordon Graham, Robin Skuce, Louis du Plessis, Suzan Thompson, Lorraine Wright, Andrew Byrne, Adrian R. Allen

**Affiliations:** University of Glasgow, Glasgow, UK; Centro de Investigacion en Alimentacion y Desarrollo A.C., Hermosillo, Sonora, Mexico; Fios Genomics, Edinburgh, UK; Department of Infectious Diseases, College of Veterinary Medicine, University of Georgia, Athens, GA, USA; Foreign, Commonwealth and Development Office, Glasgow, UK; Agri-Food and Biosciences Institute, AFBI Stormont, Belfast, UK; University of Edinburgh, Roslin Institute, Edinburgh, UK; Department of Agriculture, Environment and Rural Affairs (DAERA), Belfast, UK; Department of Zoology, University of Oxford, Oxford, UK; Department of Agriculture Food and the Marine (DAFM), Dublin, Ireland

## Abstract

**Background:** Bovine tuberculosis (bTB) is a costly, epidemiologically complex, multi-host, endemic disease. Pathogen whole genome sequencing can improve the resolution of epidemiological tracing. We genome sequenced an exceptional data set of 619 *Mycobacterium bovis* isolates from badgers and cattle in a 100km^2^ bTB ‘hotspot’. Historical molecular subtyping data permitted the targeting of an endemic pathogen lineage, whose long-term persistence provided an opportunity to study genome epidemiology in detail. To assess whether badger population genetic structure was associated with the spatial distribution of pathogen genetic diversity, we microsatellite genotyped hair samples from 769 badgers trapped in this area.

**Results:** Eight lineages of *M. bovis* were circulating in the study area, seven of which were likely non-endemic, and imported by animal movement. The endemic lineage exhibited low genetic diversity with an average inter-isolate genetic distance of 7.6 SNPs (s.d. ± 4.0), consistent with contemporary transmission. Bayesian phylogenetic methods determined an evolutionary rate of 0.30 substitutions per genome per year for this lineage, estimating its emergence 40-50 years before present, while Bayesian Skyline analysis identified significant population expansion of the endemic lineage in the 1990s and again in 2011-2012. The phylogeny revealed distinct sub-lineages, all of which contained isolates from both cattle and badger hosts, indicative of the sharing of closely related strains and inter-species transmission. However, the presence of significant badger population genetic structure was not associated with the spatial distribution of *M. bovis* genetic diversity.

**Conclusions:** Our data provided unparalleled detail on the evolutionary history of an endemic *M. bovis* lineage. Findings are consistent with ongoing interspecies transmission in the study area but suggest that badger intra-species transmission may not be a major driver of persistence in this area. In addition, the data collected permitted the tracking of incursions of novel pathogen lineages into the study area and means to determine if they were involved in disease transmission.

## 1. Introduction

### 1.1 Background

*Mycobacterium bovis* infection in cattle (*Bos taurus*) and badgers (*Meles meles*) is a persistent and costly problem for the farming industries and governments of the United Kingdom and Republic of Ireland (Allen et al. 2018). In Northern Ireland alone, the bovine tuberculosis (bTB) eradication scheme cost £44 million in 2017/2018 (Northern Ireland Audit Office, 2018). The complex epidemiology of the disease is well recognised, with the role of wildlife in transmitting infection to cattle acknowledged as an impediment to eradication (Godfray et al. 2018).

Multi-host zoonotic infections of slowly-evolving pathogens, such as the members of the *M. tuberculosis* complex (MTBC), present significant challenges to researchers who wish to use molecular epidemiological methods to understand disease transmission dynamics (Biek et al. 2015). Previously, multi-locus variable number of tandem repeats analysis (MLVA) and spoligotyping were used to characterise spatio-temporal patterns in bTB epidemiology (Kamerbeek et al. 1997; Skuce et al. 2010; Milne et al. 2019; Skuce et al. 2020). These methods have demonstrated how *M. bovis* infections typically present as a series of geographically localised micro-epidemics (Skuce et al. 2010; Skuce et al. 2020). However, MLVA and spoligotype loci, whilst extremely useful in defining the home ranges of endemic infections (Trewby, 2016a; Milne et al. 2018), evolve at rates considerably slower than the inter-host transmission rate, thereby limiting their utility for contemporary disease outbreak investigations (Meehan et al. 2018).

Whole Genome Sequencing (WGS) technologies and associated phylogenetic analytical frameworks, have helped to reveal sources of infection and to improve surveillance and control for various pathogens (Harris et al. 2013; Mellman et al. 2011; Walker et al. 2013). These phylodynamic methods have been most effectively applied to fast-evolving viral pathogens, whose mutation rates can, with dense sampling, permit inference of fine scale disease dynamics, over short time intervals (Volz et al. 2009; Biek et al. 2015).

Biek et al (2012) were the first to apply WGS and Bayesian phylogenetics to the *M. bovis* epi-system, focusing on an emerging endemic strain found in the east of Northern Ireland. Since then, a variety of phylodynamic studies using ancestral state reconstruction and structured coalescent methods, have been applied to the *M.* bovis epi-system and have helped to characterise transmission dynamics involving livestock and wildlife (Trewby et al. 2016b; Crispell et al. 2017; Salvador et al. 2019; Crispell et al. 2019, Crispell et al. 2020; Rossi et al. 2020).

Pathogen spread across landscapes is recognised to be an inherently spatial process (Biek and Real, 2010), leading to distinct patterns in pathogen genetic structure. Similarly, free ranging wildlife hosts, exhibit partitioning of their own genetic variation across landscapes (Guerrero et al. 2018), for example, isolation by distance (IBD) (Wright, 1943). An appreciation of how these types of landscape-genetic phenomena intersect can help to inform epidemiological investigations in wildlife populations (Biek and Real, 2010). A key question within localised bTB micro-epidemics is whether significant badger population structure is observed over smaller geographic scales in Ireland, and if so, whether it explains any of the partitioning of *M. bovis* genetic variation in the landscape. If badgers are playing a key role in maintaining disease in a landscape by sustaining infection among themselves and passing infection on to cattle, then one may expect that their philopatric population structure will affect the population structure of *M. bovis* – more closely related badgers, should harbour more genetically similar sequence types of *M. bovis*.

### 1.2 Objectives

In this study we sought to better understand the molecular epidemiology of this important veterinary pathogen using the highest resolution WGS data available, in a region of high pathogen prevalence. Compared to some previous studies, we undertook a systematic, sympatric sampling of both cattle and badgers. The study was conducted in a small 100 km^2^ area of Northern Ireland (Figure 1A), using data from a recently completed (2014-2018) wildlife intervention. The ‘test and vaccinate or remove’ (TVR) selective culling protocol applied to the local badger population (Arnold et al. 2021; Menzies et al. 2021) provided an important opportunity to systematically sample sympatric cattle and badger populations for *M. bovis*, and to apply WGS, phylogenetic and population genetic methods to attempt to determine the roles that both cattle and badgers play in the local disease dynamics.

**Figure 1.**
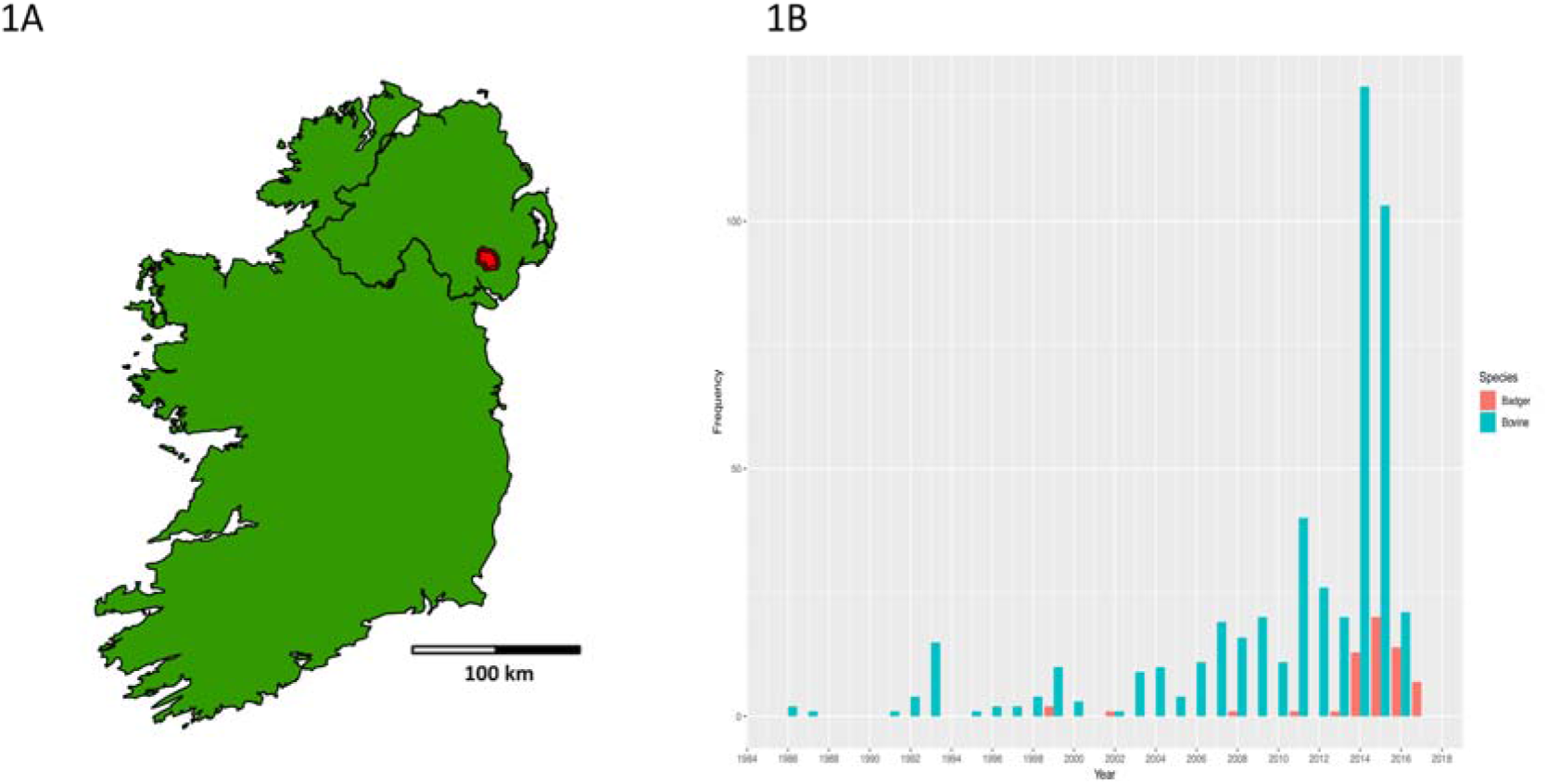
A: Map of Ireland with location of 100 km^2^ TVR cull zone highlighted in red. B: Frequency bar-plot of *Mycobacterium bovis* isolates sampled annually, from unique badgers and cattle, in the TVR zone prior to (1986-2013) and during the intervention (2014-2017).

Specifically, we sought to define the evolutionary history of an endemic pathogen lineage, its genetic diversity and the relatedness of isolates derived from cattle and badgers to broadly inform on transmission dynamics. Within this endemic lineage, we also sought to determine the effect of badger population structure on the spatial partitioning of pathogen diversity. More generally, we also sought to assess the utility of WGS in detecting the incursion of non-endemic lineages and whether they established persistent foci of infection.

## 2. Methods

### 2.1 Sampling of cattle and badgers

The TVR zone (Figure 1A), was chosen because it has a high prevalence of cattle TB (24% confirmed bTB herd prevalence over two years [2011/2012] – wider Northern Ireland average was ~6-7%), and is embedded within County Down, which also has one of the highest average badger densities (3.88 badgers per km^2^) in Northern Ireland – (Reid et al. 2012).

An initial survey was conducted in 2014 to establish sett locations (DAERA, 2018). Overnight cage trapping was used across all years between the months of May and October 2014-2018. Trapped badgers were anaesthetised, trap-side, hair sampled and tested for *M. bovis* infection using the dual path platform (DPP) serology test based on the StatPak method (Chambers et al. 2009). Tracheal aspirate was taken from all trapped badgers, whether DPP-positive or -negative and sent for bacteriological culture as described below. From 2015-2018, DPP positive badgers were humanely euthanized and DPP-negative badgers were vaccinated using injectable Bacillus Calmette Guerin (BCG), and released (Menzies et al. 2021). Culled badgers underwent post-mortem examination according to a standardised protocol (Courcier et al. 2020) and specified tissues were submitted for bacteriological culture.

All cattle in Northern Ireland are TB tested annually using the standardised (European Council, 1964) single intradermal comparable cervical tuberculin (SICCT) test (Abernethy et al. 2006). Specified tissue from all SICCT positive bTB reactor cattle in the TVR region from 2014-2017 were harvested at the time of slaughter and submitted for bacteriological culture.

To provide extra temporal depth, a selection of historical isolates (n=243) from both hosts in the TVR zone were re-cultured from the AFBI *M. bovis* Northern Ireland wide, strain archive. The temporal window for all isolates runs from 1986 to 2017. All historical badger isolates were derived from road traffic accident (RTA) post-mortems in a surveillance scheme run by the Department of Agriculture, Environment and Rural Affairs in Northern Ireland (DAERA-NI) (Courcier et al. 2018). In addition, an area directly neighbouring the study zone has a distinct lineage of *M. bovis* present, four isolates of which were detected in the study zone. A random sample of historical isolates from this neighbouring area was collected to provide additional phylogenetic context.

### 2.2 *M. bovis* culture and genomic DNA extraction

*M. bovis* isolates were initially cultured in the liquid BD BACTEC MGIT system, solid Stonebrinks and Löwenstein-Jensen media and single colonies were selected for sub-culture (Skuce et al. 2005). Isolates were heat-killed in a water bath at 80°C for 30 min. DNA was extracted using standard high salt/cationic detergent cetyl hexadeycl trimethyl ammonium bromide (CTAB) and solvent extraction protocols (Parish and Stoker, 2001; van Soolingen et al. 2002).

### 2.3 Spoligotyping and MLVA analysis

*M. bovis* isolates were genotyped by spoligotyping (Kamerbeek et al., 1997) and 8 locus MLVA using previously described methods (Skuce et al., 2010). Authoritative names for spoligotype patterns were obtained from www.mbovis.org (Smith and Upton, 2012). MLVA profiles were named using a laboratory nomenclature (Skuce et al. 2005).

### 2.4 Genome sequencing and bioinformatic analyses

Sequencing libraries were prepared using the Illumina Nextera XT method to produce inserts of approximately 500-600bp. One hundred samples were sequenced at AFBI using an Illumina MiSeq platform with Illumina V2 chemistry, producing paired-end reads of 250 bp. A further 100 samples were sequenced at the Glasgow Polyomics facility using an Illumina Miseq producing 2×300 bp paired end reads. All remaining samples were sequenced by Eurofins Scientific using an Illumina HiSeq producing 2 ×250 bp paired end reads. For quality assurance (QA) purposes, comparison of sequencing performance across all three sites was undertaken at AFBI. Specifically, we re-sequenced 15 randomly selected isolates from those sent to other institutes and compared these duplicates to the initial sequencing data.

Reads for each sample were mapped to the recently updated/annotated (Malone et al. 2017) reference genome for *M. bovis* strain AF2122/97 (GenBank accession <u>LT708304.1</u>) using the mapping-based phylogenomics pipeline, RedDog V1beta.10.3 (Edwards et al. 2016) to identify SNP variants across all isolates. Alignment and mapping were carried out in Bowtie2 v2.2.9 (Langmead et al. 2012) using the sensitive local mapping setting. SNP calling was undertaken using SAMtools and BCFtools (Li et al. 2009) using the consensus caller setting. A minimum depth of 10x was set for SNP calling. The average coverage failure filter, average depth filter and average mapping failure filters were set at 98%, 10x, and 75% respectively. Transposable elements, repeat regions and the PE/PPE regions as defined in the Genbank annotation, were excluded from SNP calling using the parseSNP tool in the RedDog pipeline (Edwards et al. 2016).

Some isolates were sequenced by Glasgow Polyomics, others were sequenced by Eurofins laboratories. 15 randomly selected isolates were re-sequenced by AFBI. No SNP distance was observed between duplicates and the previously sequenced isolates.

### 2.5 Badger microsatellite genotyping

Nuclear DNA extracted from all badger hair samples was genotyped at 14 microsatellite loci using methods previously described (Guerrero et al. 2018). We re-profiled 5% of DNA samples as a quality assurance measure.

### 2.6 Testing for IBD in badger population

Using the package ‘PopGenReport’ (Adamack and Gruber, 2014) in R (R Development Core Team, 2020), we constructed a microsatellite distance matrix (Smouse and Peakall method) for all unique badgers captured in the study zone, and for the subset of badgers that produced *M. bovis* cultures from the endemic lineage (see below), to ensure we had sufficient power to detect IBD in sub-populations. For both datasets, we then constructed inter-animal Euclidean distance matrices using the R package, ‘Geosphere’ (Hijmans et al. 2019). We then performed a Mantel test, with 10,000 repetitions, for IBD using the package ‘ade4’ (Dray and Dufour, 2007) in R. For the larger meta-population, Mantel tests and linear regressions were carried out for each capture year, with an analysis of covariance carried out to compare the slopes of each genetic distance vs euclidean distance relationship.

### 2.7 Phylogenetic analyses

The most appropriate nucleotide substitution model for our phylogenetic analyses of the FASTA alignments of informative SNPs was assessed using the ‘modelTest’ function of the package ‘Phangorn’ (Schliep, 2011) in R. Specifically, the fit of the General Time Reversible (GTR), Jukes Cantor, and Hasegawa Kishino Yano (HKY) models to the data were assessed. The nucleotide substitution model with the lowest AIC (GTR – AIC: 26860.39) was used to build a maximum likelihood phylogeny using RAxML v8 (Stamatkis, 2014). The rapid bootstrapping search method in RAxML was selected and stopped after 400 replicates. The phylogeny was visualized and assessed in FigTree v1.4.4 (Rambaut, 2018) and the ape package (Paradis and Schliep, 2018) in R. Final figures of maximum likelihood trees were produced using ggtree (Yu, 2020). The presence/absence of a temporal signal in the phylogenetic data of the endemic clade was assessed using the program TempEst v1.5.1 (Rambaut *et al*. 2016). After selecting the reference strain (AF2122/97) as the bestfitting, outgroup root isolate, which maximises the temporal signal, the root to tip divergence model was fitted using the residual mean squared method.

To test the significance of the temporal signal in our dataset, we randomised the tip dates for the 302 chosen isolates from the endemic clade, in ten replicate analyses, as *per* Firth et al (2010). Tip dates were randomised using the ‘Tipdatingbeast’ package (Rieux and Khatchikian, 2017) in R. The original dataset and each replicate were subjected to a simplified BEAST 2 (Bayesian Evolutionary Analysis by Sampling Trees - Bouckaert et al. 2014), constant population, coalescent analysis, using a relaxed clock model, the GTR nucleotide substitution model. A chain length of 100,000,000 MCMC steps was used for the three non-randomised tip date runs, with a longer 300,000,000 MCMC steps used for all ten randomised runs. 10% of MCMC steps were discarded as burn-in. After checking convergence in Tracer 1.7.1 (Rambaut et al. 2018) – (ESS > 200 for all parameters), the median substitution rate for each replicate was compared to those of the logcombined non-randomised data set.

To determine the past demographics of the endemic *M. bovis* lineage in the study region, we also performed a Bayesian Skyline analysis in BEAST 2 (Drummond et al. 2005). As above, we used a relaxed lognormal clock and the GTR nucleotide substitution model. We set the model to assess historical effective population size (N_e_) over 4 dimensions. A chain length of 200,000,000 MCMC steps was used for three replicates. 10% of steps were discarded as burn in and replicate log and tree files were then combined in logcombiner and analysed in Tracer 1.7.1.

Also within the endemic *M. bovis* clade, we ran a Mantel test to determine if closely related sequence types exhibited any geographic localization / clustering within the study area. We constructed an inter-isolate SNP distance matrix using the ‘Biostrings’ package (Pagés et al. 2021), and an inter isolate Euclidean distance matrix in ‘Geosphere’ (Hijmans et al. 2019) in R. The Mantel test was run for 10,000 iterations in the ‘ade4’ package in R as described above.

### 2.8 Assessing effect of badger population structure on *M. bovis* spatial partitioning

We sought to determine if *M. bovis* inter-isolate SNP distance is associated with pairwise Euclidean distance between trapped badger locations, pairwise difference in time of *M. bovis* isolation and pairwise host genetic distance.For *M. bovis* genome sequences derived from badgers infected with the endemic lineage, we constructed two inter-isolate distance matrices: (i) SNP distance using the R package ‘ape’ (Paradis and Schliep, 2019); (ii) time of isolation difference using the ‘dist’ function in R.

Using these *M. bovis* distance matrices and the two already produced to assess IBD in the infected badgers (see above), we performed a multiple regression on distance matrices (MRM) analysis using the R package ‘ecodist’ (Goslee and Urban, 2007) with 10000 repetitions.

## 3. Results

### 3.1 Sampling of cattle and badgers

A total of 642 *M. bovis* isolates were used in this study, of which 611 were collected from badgers and cattle in the TVR zone, and 31 from a neighbouring region. Of the 642 isolates, 15 were sequencing QA duplicate controls as discussed in section 2.4. In addition, we re-sequenced the *M. bovis* reference sample, AF2122/97 as an internal control. Of the 642 survey isolates, 399 (282 cattle; 117 badger) were sampled contemporaneously in the TVR zone during the project (2014-2017). A further 242 historical isolates (232 cattle; 10 badger=10) from 1986-2013 were available from archived cultures from the zone and its environs. Cattle isolates across all years were single isolates per animal from 185 herds. Multiple isolates (n=86) were cultured from 24 individual badgers, with single isolates derived from a further 36 badgers. In total, between 1986 and 2017, we collected *M. bovis* isolates from 60 unique badgers and 483 unique cattle (Figure 1B). Full details of sample locations, year of isolation and species of origin are given in Supplementary Data 1.

### 3.2 Spoligotyping and MLVA analysis

22 MLVA types and 6 spoligotypes were observed for the 642 isolates. From prior analyses (Skuce et al. 2010), the spoligotype and MLVA genotypes could be grouped into eight related ‘strain families’. Each is dominated by a probable founder genotype (source of the family name) with related daughter strains varying by spoligotype and / or MLVA polymorphism. Numbers of isolates per strain family are shown in Table 1A. The MLVA 6, spoligotype 263 family (6.263) was considered to be endemic in the region and accounted for most of the observed isolates (Skuce et al. 2010; Skuce et al. 2020). The remaining seven strain families (1.140, 2.142, 3.140, 4.140, 5.140, 19.140 and 20.131) were not likely to be endemic in the TVR cull zone as each has a home range elsewhere in Northern Ireland (Skuce et al. 2010; Trewby, 2016a). Of the 36 isolates from strain family 20.131, 32 were collected from a region neighbouring the TVR zone, 4 badger isolates were found within the zone. Full details of isolate MLVA genotypes and spoligotypes are supplied in Supplementary Data 1.

**Table 1.**
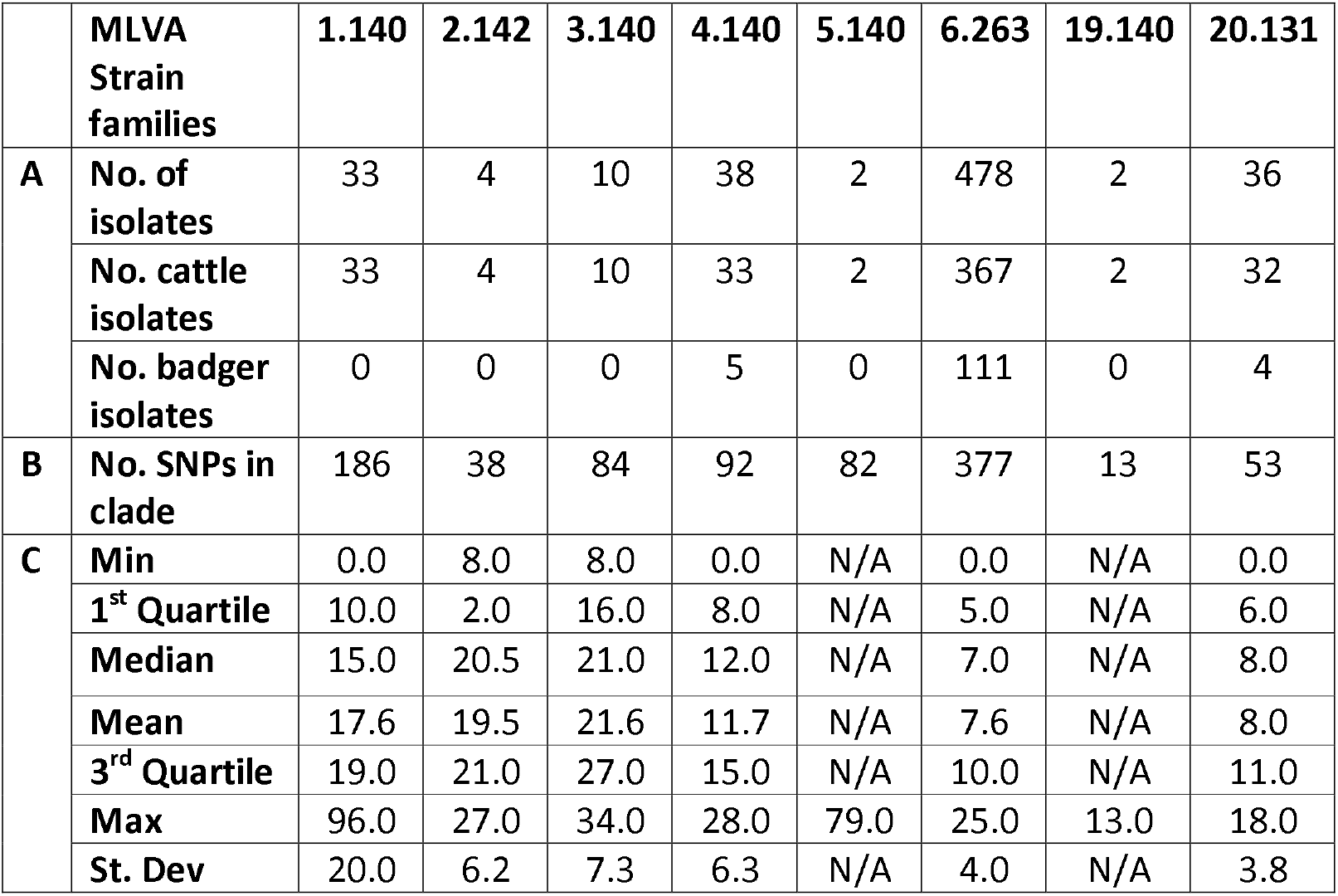
A. number of isolates per strain family sampled in the study area with breakdown of number of isolates per host species. B. number of SNPs detected in strain family clades, and C. pairwise SNP distance statistics for each of the eight major lineages of *M. bovis* found in the TVR zone.

### 3.3 Sequencing, bioinformatic analyses and Quality Assurance

The RedDog pipeline was used to process the isolates. 24 (22 cattle and 2 badgers) failed the sequencing QA filters (98% coverage filter for the reference genome) and were excluded. The remaining 618 survey isolates plus AF2122/97 control passed all QA filters. Detailed QA meta-data for all 619 isolates are given in Supplementary Data 2. Forward and reverse reads for all QA passing isolates are deposited at the National Centre for Biotechnology Information (NCBI) Sequence Read Archive (SRA), bio-project XXXXXXXX. Accession numbers for all reads are given in Supplementary Data 2.

From the 619 isolates with good quality sequence reads, 1562 SNPs passed QA calling rules and were used to conduct phylogenetic analyses. Details of all SNPs passing QA, and their location in the reference sequence are given in Supplementary Data 3.

The AF2122/97 control exhibited a 3 SNP distance from the reference sequence (Genbank <u>LT708304.1</u>) likely due to accrual of mutations after culture passages at AFBI.

### 3.4 Badger microsatellite genotyping and IBD analyses

769 unique badgers were captured, location recorded, sampled and successfully genotyped between 2014 and 2018. Random 5% re-genotyping for QA purposes produced identical allele calls. Microsatellite profiles, capture locations and date of capture can be found in Supplementary Data 4. Summary population genetic statistics for all animals, per year are collated in Supplementary Table S1.

Samples from 45 badgers produced positive *M. bovis* cultures. Spoligotyping and MLVA placed them in the major endemic lineage in the study zone. Capture locations for the 45 endemic strain positive badgers are illustrated in Supplementary Figure S1A.

Across all years, the badger population exhibited consistent levels of IBD, as indicated by significant Mantel tests (r=0.11-0.17, p<0.05) (Table 2). The slopes of the relationships between genetic distance and Euclidean distance were very similar. Small, significant differences were observed in the ANCOVA (Supplementary Figure S2 and Table 2), mainly due to the increased slope of the IBD relationships years 2015, 2016, and 2018, consistent with badger genetic differentiation being observed over shorter distances.

**Table 2.** Badger meta population isolation by distance (IBD) relationship for all sampling years. * =2015 significantly different than slopes for years 2014, 2016 and 2017. ** = 2016 slope significantly different than slopes for years 2014 and 2015. ***=2018 slope significantly different than slopes for years 2014 and 2017.

| Cohort genotyped | No. animals | Mantel test Pearson coefficient r | Mantel p value | Linear model beta | Linear model p value | R<sup>2</sup> | 1 unit differentiation per x km distance |
| --- | --- | --- | --- | --- | --- | --- | --- |
| **2014** | 273 | 0.11 | 0.0001 | $1.6 \times 10^{-4}$ | $< 2 \times 10^{-16}$ | 0.012 | 6.25 km |
| **2015** | 152 | 0.16 | 0.0001 | $2.1 \times 10^{-4}$ | $< 2 \times 10^{-16*}$ | 0.024 | 4.74 km |
| **2016** | 97 | 0.17 | 0.0001 | $2.6 \times 10^{-4}$ | $< 2 \times 10^{-16**}$ | 0.030 | 3.84 km |
| **2017** | 113 | 0.11 | 0.0001 | $1.5 \times 10^{-4}$ | $< 2 \times 10^{-16}$ | 0.011 | 6.67 km |
| **2018** | 134 | 0.13 | 0.0004 | $1.7 \times 10^{-4}$ | $< 2 \times 10^{-16***}$ | 0.017 | 5.88 km |
| **TB +ve** | 45 | 0.16 | 0.0024 | $3.1 \times 10^{-4}$ | $1.1 \times 10^{-7}$ | 0.030 | 3.22 km |

The 45 *M. bovis* culture positive badgers also exhibited significant IBD (Table 2) similar to that of the larger study population.

### 3.5 Preliminary phylogenetic analyses

#### 3.5.1. All isolates

The Maximum Likelihood tree constructed in RAxML for all 619 sequenced isolates is shown in Figure 2. The phylogeny was rooted using the 20.131 strain family as an out-group, as these isolates are known to derive from an older common ancestor than other extant strains (Allen et al. 2013). Eight major lineages, each with high bootstrap support, were observed in the phylogeny. The eight strain families defined previously by MLVA and spoligotyping were monophyletic and in perfect concordance with the SNP based tree topology. We determined the within-lineage diversity, as defined by total number of SNPs recorded, for each of the eight major lineages observed (Table 1B). Additionally, from distance matrices generated during phylogenetic analyses, pairwise, inter-isolate SNP distance, summary statistics were computed within all eight major lineages (Table 1C).

**Figure 2.**
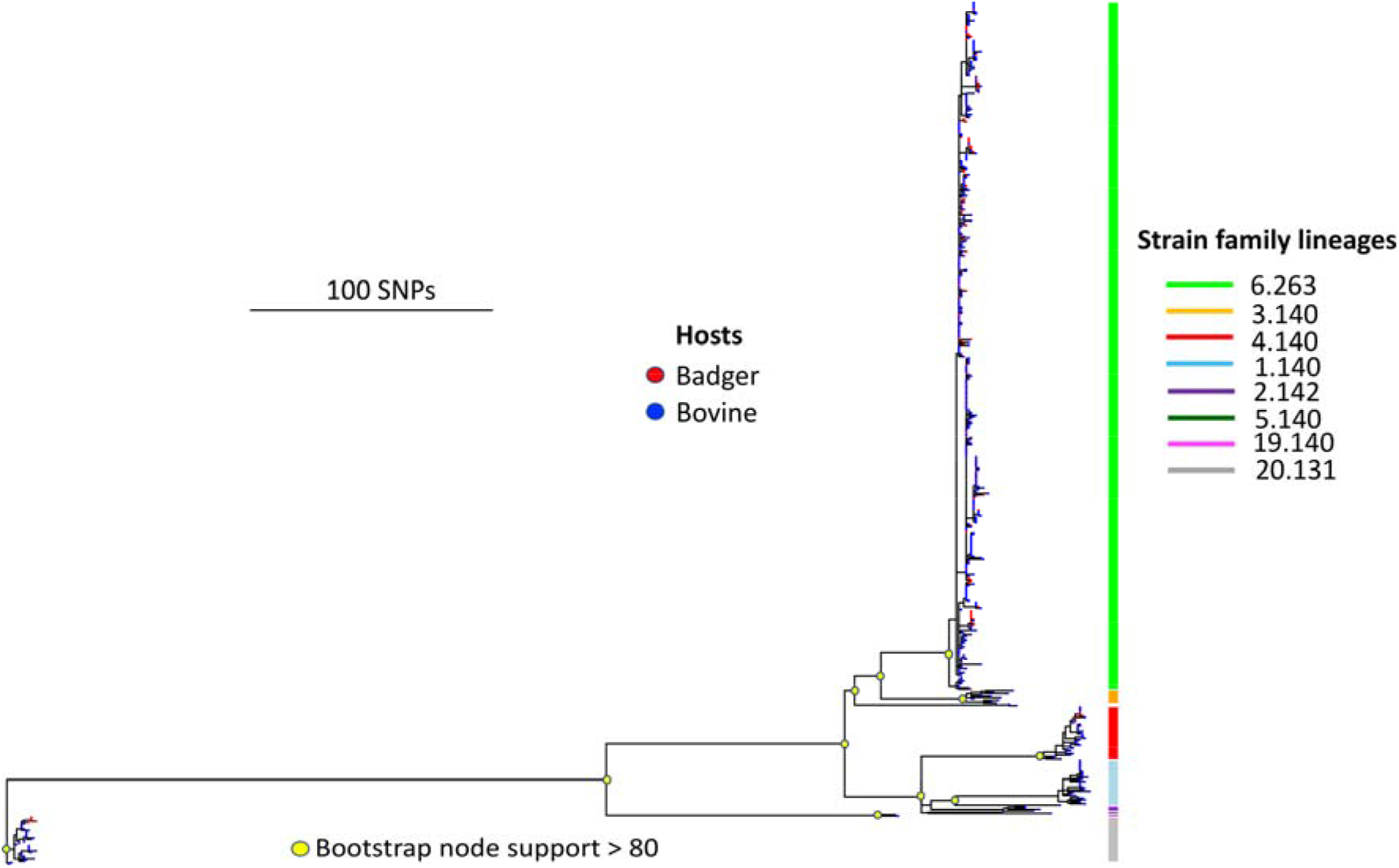
1562 SNP Maximum Likelihood phylogeny of all 619 isolates that passed sequencing QA.

#### 3.5.2 Endemic lineage – 6.263

The endemic major lineage of 6.263 presented the best opportunity to investigate *M. bovis* transmission dynamics among cattle and badgers in the TVR zone. 6.263 has been consistently associated with the study area for over two decades in local cattle and badgers, unlike lineages whose home range is elsewhere in Northern Ireland (Skuce et al. 2020). A higher resolution SNP phylogeny of 6.263 is shown in Figure 3, using the subset of 302 isolates described above. Cattle and badger isolates were observed in all sub-lineages of the endemic lineage, with no sub-lineages made up of isolates exclusively from a single host species. Major sub-lineages had good bootstrap support (>90). A bar-plot showing frequency of isolates per host species taken from the endemic clade over the period 1986-2017, is presented in Supplementary Figure S3. A smaller maximum likelihood phylogeny of the 45 endemic clade badgers is presented in Supplementary Figure S1B.

**Figure 3.**
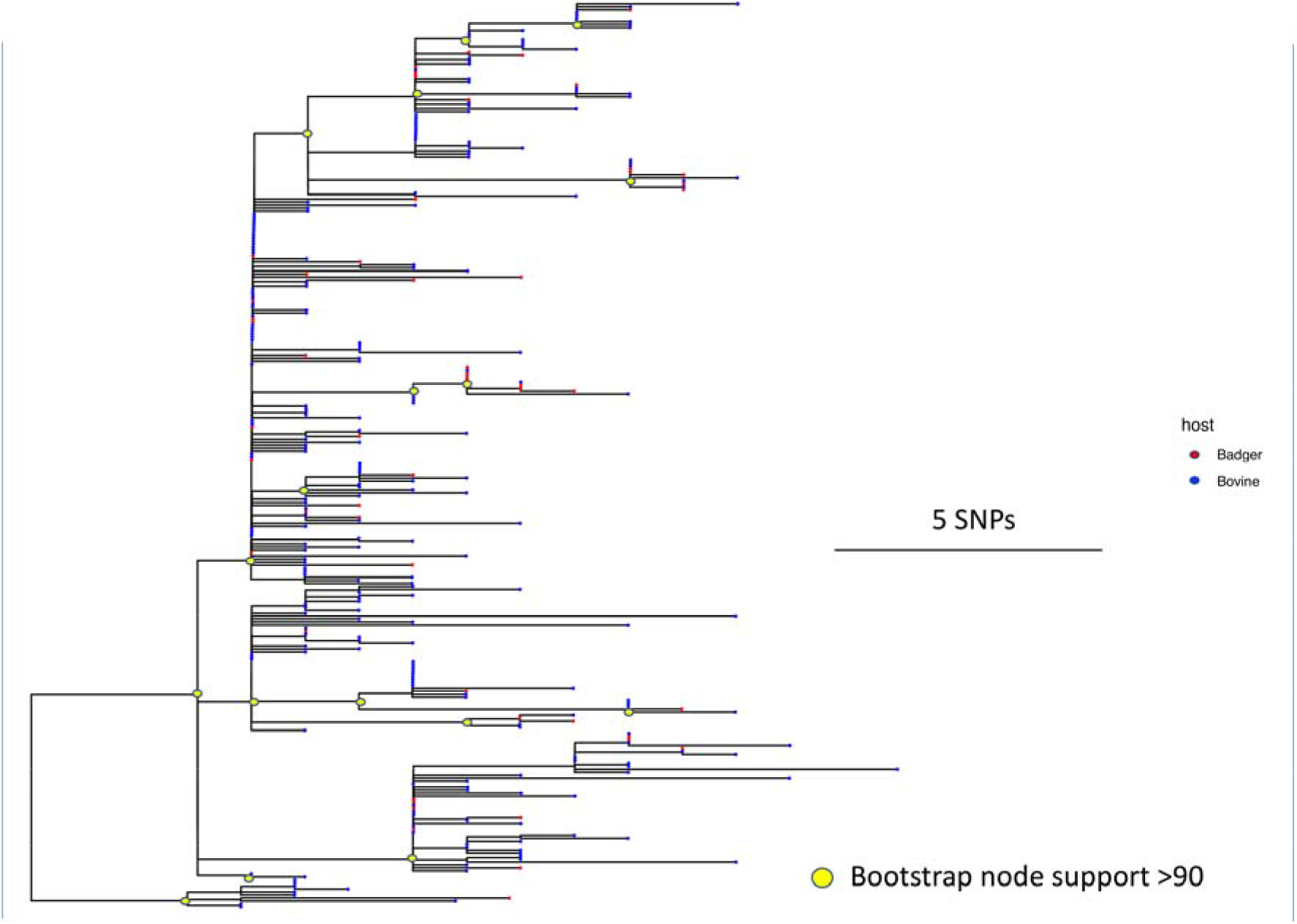
377 SNP Maximum Likelihood phylogeny of 302 isolates from the 6.263 endemic lineage in the TVR zone. Tree rooted to AF2122/97 reference, but reference removed for aid of visualisation.

Badger isolates were predominantly found in the endemic 6.263 lineage. No badger isolates were found in the 1.140, 2.142, 3.140, 5.140 and 19.140 lineages. Four isolates, from two unique badgers, sampled in the TVR zone, were found in the 20.131 lineage, whilst five isolates from four unique badgers were found in the 4.140 lineage (Supplementary Figures S4 and S5). A full breakdown of number of isolates from each species in each lineage is shown in Table 1A.

The n=302 endemic lineage phylogeny analysed in TempEst, rooted against the AF2122/97 reference genome, exhibited a positive correlation between genetic divergence (root to tip distance) and sampling time, with moderate evidence of temporal signal (R^2^=0.25; p<0.001), and a conservative clock rate of approximately 0.22 substitutions per genome, per year - see Supplementary Figure S6A. All ten tip date randomised replicate datasets run using the simple coalescent model, exhibited similar substitution rates, all of which were considerably lower than the substitution rate inferred on the non-randomised data set (0.30 substitutions per genome, per year 95% HPD: 0.24-0.37) and exhibited no overlap in 95% highest posterior density (HPD) (Supplementary figure S6B).

The Bayesian Skyline analysis indicated a substitution rate that was very similar to that observed in the non-randomised simple coalescent model described above (0.35 substitutions per genome, per year 95% HPD:0.28-0.41-Supplementary Figure S6B). It also indicated that after the origin of the 6.263 clade in the 1970s-1980s, the effective population size substantially expanded in the early to mid 1990s, and again in 2011-2012 prior to the TVR trial beginning (Figure 4).

**Figure 4.**
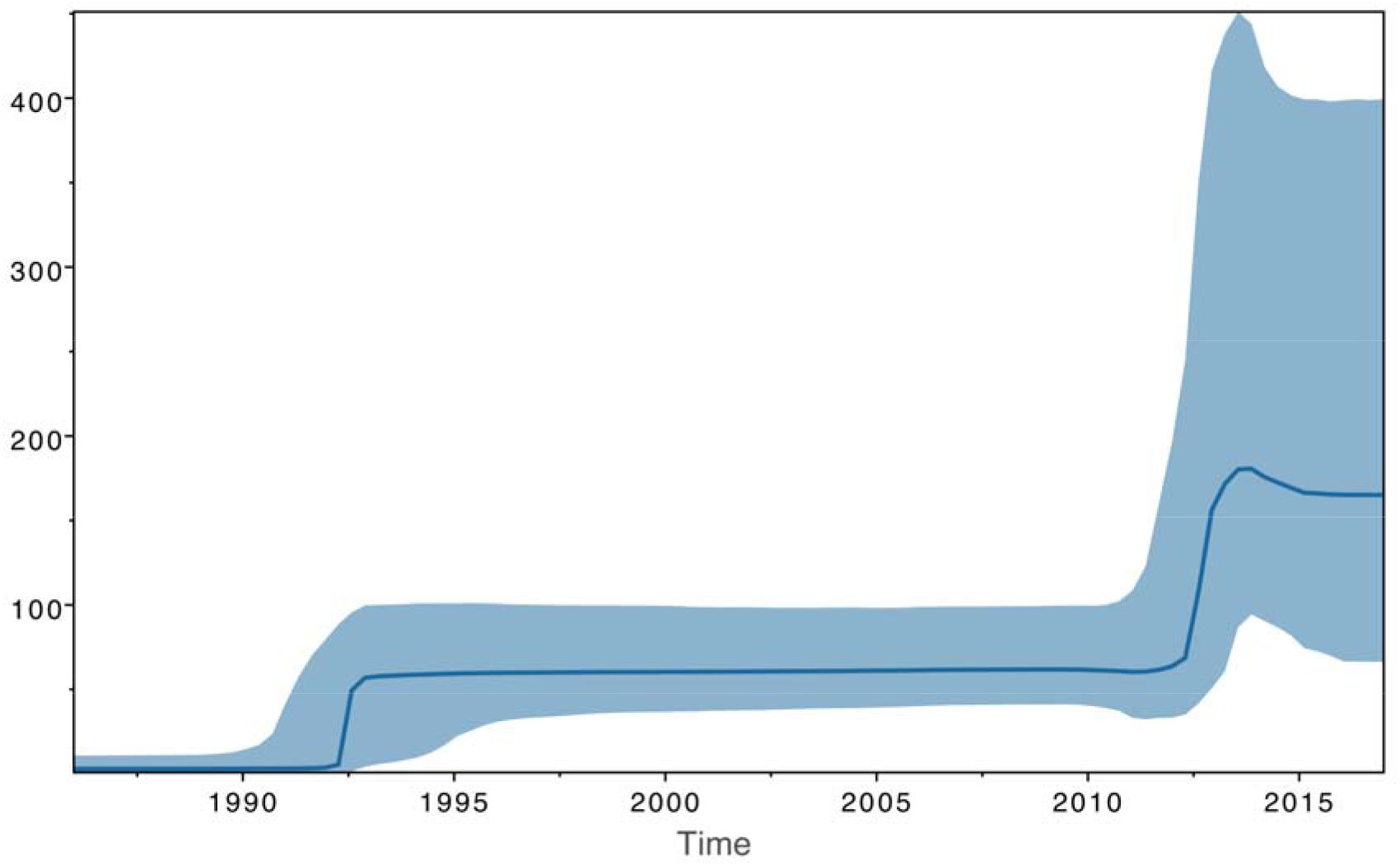
Bayesian Skyline plot of the 6.263 endemic lineage.

The Mantel test conducted for the endemic clade indicated that there was no significant spatial autocorrelation for sequence types (correlation coefficient = 0.04, p=0.157) – consistent with there being no significant clustering of closely related sequence types within the study area – see Supplementary Figure S7. An illustration of the spatial distribution of sub-clades within the endemic 6.263 lineage, and their substantial overlap, is provided in Supplementary Figure S8.

### 3.6 Effect of badger population structure on *M. bovis* spatial partitioning

From the full model in the MRM analysis, modeling *M. bovis* genetic distance (SNP-based) as a function of inter badger genetic distance, inter badger Euclidean distance and inter *M. bovis* time of isolation difference, we observed that only inter-badger Euclidean distance was significantly associated with *M. bovis* genetic distance (p = 0.04). However, the overall fit of the model was non-significant (F-test p >0.05). Badger microsatellite derived genetic relatedness was therefore not associated with *M. bovis* SNP-derived genetic differentiation. A full summary of MRM findings is presented in Supplementary Table S2.

## 4. Discussion

### 4.1 General observations on WGS for enhanced molecular epidemiology

Our data are consistent with the findings that WGS provides unparalleled resolution for epidemiological investigations of zoonotic disease (Kao et al. 2014; Kao et al. 2016) – indexing additional pathogen variation to stratify isolates that are homogeneous according to the classical tuberculosis molecular epidemiological tools of spoligotyping and MLVA (Skuce et al. 2010; Skuce et al. 2020).

Regarding the latter tools, we find perfect congruence between spoligotype and MLVA data and the basal nodes defining major lineages of the phylogeny for all isolates from the TVR region (Figure 2). This concordance is testament to the clonality of *M. bovis* (Smith et al. 2006; Allen et al. 2013) and is indicative that existing databases of classical molecular markers, can be used to target ‘hotspots’ of persistent, endemic infection for closer investigation using WGS.

Contemporary transmission of slowly evolving pathogens, such as members of the *Mycobacterium tuberculosis complex* (MTBC), is typically characterised by little within clade diversity and resulting reduced SNP distances / phylogenetic branch lengths between isolates (Meehan et al. 2018). Previous studies have suggested various minimal SNP distance thresholds, for the definition of contemporaneous, epidemiologically linked isolates. Five and twelve SNP distances have been proposed to be consistent with such transmission clusters in *M. tuberculosis* outbreaks in the United Kingdom (Walker et al. 2013; Walker et al. 2014), whilst ten SNP thresholds have been proposed in other studies with *M. tuberculosis* and *M. bovis* (Bryant et al. 2013; Roetzer et al. 2013; Yang et al. 2017; Jajou et al., 2018; Crispell et al. 2019). Meehan et al (2018) have observed that in *M. tuberculosis*, SNP distances between one and five can represent transmission events up to 10 years apart. Given that MTBC evolution appears to consistently involve relaxed molecular clock like behaviour across lineages (Crispell et al. 2017; Crispell et al. 2019; Menardo et al. 2019), it is perhaps not surprising that selecting a definitive threshold for contemporary transmission is difficult, and as a result, those set can appear arbitrary. In addition to the issues with relative clock like behaviour of different *M. bovis* lineages, intensity of sample collection and representation of multiple time points can be crucial to establishing robust substitution rates (Kao et al. 2016). The general rule of thumb remains however - the shorter the SNP distance between isolates, the more likely they are more closely epidemiologically linked. In this study, the lineage we know to be endemic in the study area from years of MLVA surveillance, exhibits the shortest average pairwise SNP distance between isolates (7.6 s.d. ±4.0 – see Table 1), which is likely indicative of contemporary transmission in the region, and compares favourably to the thresholds discussed above.

The advantage that WGS will provide in disease tracing compared to historical molecular epidemiology methods in the *M. bovis* epi-system, is that outbreaks can be traced back to higher resolution, WGS defined lineages and sequence types, found in more precise locations than those defined by genetically homogeneous, MLVA home ranges that can cover substantial geographical areas (Skuce et al. 2010; Skuce et al 2020). A recent and pertinent example of this, is an outbreak from Cumbria in northwest England, which genome sequencing revealed was linked to the outbreak area under study here (Rossi et al. 2020).

### 4.2 Endemic 6.263 lineage

Our Bayesian phylogenetic analysis suggests that the endemic 6.263 lineage has been present in the TVR zone since the 1960s-1980s. The rate of molecular evolution of 0.30-0.35 substitutions per genome per year is consistent with previous *M. bovis* phylodynamic studies – see Supplementary Table S3. The presence of bacterial isolates from both cattle and badgers throughout all sub-lineages of the endemic 6.263 lineage, and their close genetic relatedness, indicates likely bi-directional transmission between both species. The Bayesian Skyline analysis of the endemic clade was indicative that the population size of the pathogen expanded in the early to mid 1990s, and again substantially prior to the beginning of the sett survey and TVR protocol described here from 2014-2017 – a finding congruent with the increase in herd prevalence observed in Northern Ireland over the same time-period (Robinson, 2015), and more particularly with the higher than average 24% herd prevalence observed in the study zone from 2011/2012. The magnitude of the demographic expansion observed violates the assumptions of constant population size that structured coalescent models require to accurately infer transition rates between different states (DeMaio et al. 2015; Müller et al. 2018), therefore it was not possible to undertake robust analyses with this dataset. Exploratory analyses using different structured coalescent methods (data not shown) yielded contradictory outcomes dependent on assumptions made about pathogen effective population sizes within hosts. In cases wherein substantial changes to population size have occurred, structured coalescent methods can then attempt to explain variation in terms of population structure, resulting in some within state effective population sizes to be artificially low, and some transition rates to be artificially high (du Plessis et al 2020).

More generally, the clonal expansion of *M. bovis* in this period, after initial success of the test and slaughter protocol of the 1950s and 1960s in other regions across the UK has been noted before, and suggested to potentially result from the adaptive spread of a favourable mutation or invasion of a new territory (Smith et al. 2003; Smith et al. 2006). Future work should seek to determine if similar demographic expansions are observed in other lineages and to search for candidate functional polymorphisms and signatures of selection that could underpin any potential pathogen phenotype of interest to control schemes and the development of diagnostic and vaccine tools.

The lack of spatial clustering of closely related sequence types from the entire endemic clade is an intriguing finding given that a general feature of *M. bovis* molecular epidemiology to date in the Northern Irish landscape has been association of molecular types with specific regions (Skuce et al. 2020). It may be however that while the genomic data is sufficiently high resolution to detect such an association, the location data for cattle may not be so useful. The data used are geolocations of the major farm holdings and may not necessarily be representative of the locations at which cattle have been exposed to *M. bovis*, owing to factors of farm fragmentation and the common practice of land rental distant from farm buildings (concacre), which is a feature of Northern Irish agriculture (Allen et al. 2015). This deficiency in the recording of cattle data is in contrast to the endemic *M. bovis* positive badger data which have an exact capture location, and which exhibited some evidence of spatial clustering as per the MRM analysis described above. Future use of WGS data for precision molecular epidemiology should aim to use better quality geolocation data for cattle derived isolates. Weighted centroids of pasture and farm buildings may be a more useful metric.

Badger genetic population structure, whilst remaining stable over the intervention period, had no association with how *M. bovis* genetic diversity was spatially distributed. It is possible that this lack of association is due to factors other than reduced badger to badger intraspecies transmission dynamics. The endemic lineage, if it were a relatively recent incursion, may have had little time to establish foci of persistent infection in badgers, and diffuse across the landscape through philopatric contact networks. This lineage has, however, been present in the region for ~33-48 years, providing ample time for establishment to occur. Alternatively, perturbation of the badger population, and associated dispersal arising through the application of culling, even at a small-scale (Bielby et al. 2014), may have served to obscure any association between pathogen and host population structures. The relative stability of the IBD relationship we observe, and the fact that during the culling period of 2015-2018, genetic differentiation occurs over shorter distances than the survey-only year of 2014, are consistent with no perturbation signal. GPS collar data from badgers from this same region have also suggested badger ranging did not increase significantly in cull years compared to survey (O’Hagan et al. 2021). Given these observations, we postulate that our badger genetic data may be indicative that intra-species badger transmission of *M. bovis* may not be a major driver of endemic disease persistence in the study area.

### 4.3 Monitoring of non-endemic lineages

Sequence data are useful for identifying probable incursions of non-endemic disease lineages into new areas, and for potentially determining if the incursion results in contemporary transmission and persistence. Intra-lineage, inter-isolate SNP distances greater than that observed for the endemic 6.263 lineage, are possibly indicative of a lack of contemporary transmission, likely associated with lineages which are non-endemic in the study region as per previous observations from Northern Ireland wide molecular epidemiological surveillance (Skuce et al. 2010; Skuce et al. 2020). This certainly appears to hold true for lineages 1.140, 2.142 and 3.140, which exhibit average pairwise inter-isolate distances of between 17.6 and 21.6 SNPs. Lineages 5.140 and 19.140 are only represented by two isolates each, but for both lineages, inter-isolate distances are again observed to be larger than the endemic lineage at 79 SNPs and 13 SNPs respectively. Given this observation, and the fact that we know these five lineages are outside of their MLVA defined home ranges (Skuce et al. 2020), it seems probable that they could have arrived in the study area through multiple, long distance, cattle movements. It is noteworthy, but anecdotal, that these lineages are comprised solely of isolates from cattle, suggestive perhaps that the lack of contemporary transmission for these incursive strains has resulted in no infection reaching the wildlife population. However, with deficiencies in sampling, badger ‘trappability’ and TB test diagnostics as discussed in the supplementary materials, one cannot be definitive that a non-endemic, visiting pathogen lineage has not established a focus of infection in the study area. Only continued surveillance over a wider temporal window could assess that. The establishment of long-term genome-based surveillance systems could in the future help to inform on successful incursions (Gardy and Loman, 2018).

The 20.131 lineage exhibits mean inter-isolate SNP distances which are comparable to that of the endemic 6.263 lineage (Table 1). However, only four isolates of this lineage, from two badgers (See Supplementary Figure S4), were found within the study area, with the majority sampled from a neighbouring region in which this strain has a focus of infection. The short inter-isolate distances observed are therefore more likely to be consistent with contemporary transmission in the neighbouring region. It is noteworthy however that the two badgers sampled for this lineage and found in the study zone were found in isolation, with no associated, contemporary, study zone cattle isolates. It could be that badgers have dispersed from the neighbouring region into the study zone, carrying infection with them, which has yet to appear in the cattle population. However, it is also possible that an undetected reservoir of cattle may have entered the study zone and transmitted infection to local badgers. Alternatively, an undetected reservoir not picked up by sampling could be residing within the zone. With so few isolates of the 20.131 lineage from within the study area, and the previously mentioned biases in sampling, it is impossible to be definitive. Again, detailed, longitudinal, genome facilitated surveillance would perhaps be able to inform more fully on this incursive lineage in this region.

Interestingly, one of the historically non-endemic lineages (4.140) (Skuce et al. 2020), does appear to be persisting in the TVR zone, with multiple badger-sourced isolates observed to exhibit shorter inter-isolate SNP distances (0-1) between each-other and cattle from the area (Supplementary Figure S5), consistent with more contemporary transmission events. The latter observation does highlight the usefulness of this WGS based approach for on-going surveillance, and for detecting incursions that establish new foci in new regions. It seems most probable from this case, that new foci of non-endemic lineages are introduced by cattle movements, with subsequent spill-over to badgers.

## Conclusions

Our data serve to further illustrate the utility of WGS for bovine tuberculosis molecular epidemiology, indicating ongoing transmission of infection in an endemically affected region. Our novel use of badger population genetic data alongside pathogen genomic data has also helped to demonstrate that in this particular region, this philopatric species’ stable population structure is not associated with *M. bovis* population structure, which may be indicative that badgers are not playing a major role in disease persistence. The latter may not be the case in other regions however and will require empirical assessment.

Our findings also illustrate the importance of assessing the past and recent demography of endemic lineages. Significant expansion and contraction of pathogen population size can limit the utilisation of phylodynamic tools to more closely inform on inter host transition dynamics.

More generally, the high-resolution data presented herein, serve to shrink the home range over which disease tracing investigations need to take place in comparison to MLVA based methods, thereby improving precision and inferences on likely infection sources. Developing national databases for other common, endemic lineages associated with other locations will substantially modernise field epidemiology and control schemes, enabling detection of incursions of novel sequence types into regions that are not their traditional home range (Rossi et al. 2020), and determining if such incursions have resulted in ongoing transmission between cattle and wildlife, thereby establishing novel foci of infection. A key requirement to make best use of data in the ways described will be improving the precision with which cattle geo-location data are collected.

Finally, in addition to the molecular epidemiological information that a national database of WGS data would provide, the sequence data collected would also be a useful resource for the study of evolutionary questions pertaining to possible adaptive phenotypes of *M. bovis* that may be influencing the success of the eradication scheme across Britain and Ireland.

**Table 3.** Summary statistics for log-combined simple Bayesian coalescent and Bayesian Skyline models. All estimates are median and 95% highest posterior density (HPD). Substitution rate is expressed as sites per genome, per year. Time to most recent common ancestor (tMRCA) for the endemic 6.263 clade is expressed in years prior to the last sampling date, 2017.

|  | Substitution rate | Time to most recent common ancestor<br>(tMRCA) - yrs |
| --- | --- | --- |
| Simple<br>Bayesian<br>coalescent | 0.30 (0.24-0.37) | 48.87 (33.88-82.56) |
| Bayesian<br>Skyline | 0.35 (0.28-0.41) | 33.22 (31.00-37.81) |

## Acknowledgments

Assel Akhmetova is supported by a Bolashak International Scholarship. This work was funded by the Department of Agriculture, Environment and Rural Affairs for Northern Ireland (DAERA-NI) through its ‘Evidence and Innovation’ programme – project no. 15/3/07. Additional funding was provided by the UK’s Biotechnology and Biological Sciences Research Council (BBSRC) – grant numbers BB/P0105598 and BB/M01262X. The funders had no role in study design, analysis or decision to publish. The authors wish to recognise Shane Collins and Carl McCormick and their team for supervising all field work and AFBI Disease Surveillance and Bacteriology teams who provided excellent post-mortem examinations and mycobacteriology support. Thanks to Dr Dez Delahay of the Animal and Plant Health Agency (APHA) in GB for advice on surveying local badger populations and trapping.

## Research Ethics

All badger field work was carried out under licences issued by the Northern Ireland Environment Agency. All scientific procedures performed on badgers were conducted according to the guidelines of the Animals Scientific Procedures Act (ASPA - Licence 2767) overseen by the Department of Health for Northern Ireland.

## Conflict of interest

The authors declare no conflict of interest in the production of this work.

## Supplementary Files

R scripts and BEAST xml run files used in the performance of this work are curated at the following GitHub repository:

https://github.com/AdrianAllen1977/Genome-epidemiology-of-Mycobacterium-bovis-infection-in-contemporaneous-sympatric-badger-and-cattle

## Notes

### Competing Interest Statement

The authors have declared no competing interest.

### Summary of Updates

After careful consideration and re-analysis, we, the authors, believe the structured coalescent analysis methods used to assess inter- and intra-species transmission were not suitable for the specific population under study. The population in question underwent a substantial increase in effective population size, which violates one of the core assumptions of structured coalescent models, and can lead to spurious results - we discuss this in this revised version. We have removed all estimates of transitions etc derived from these methods in this version of the manuscript and are actively exploring other ways of assessing inter-and intra-species transmission.

