## Supplementary Figures for "Genomic epidemiology of *Mycobacterium bovis* infection in sympatric badger and cattle populations in Northern Ireland"

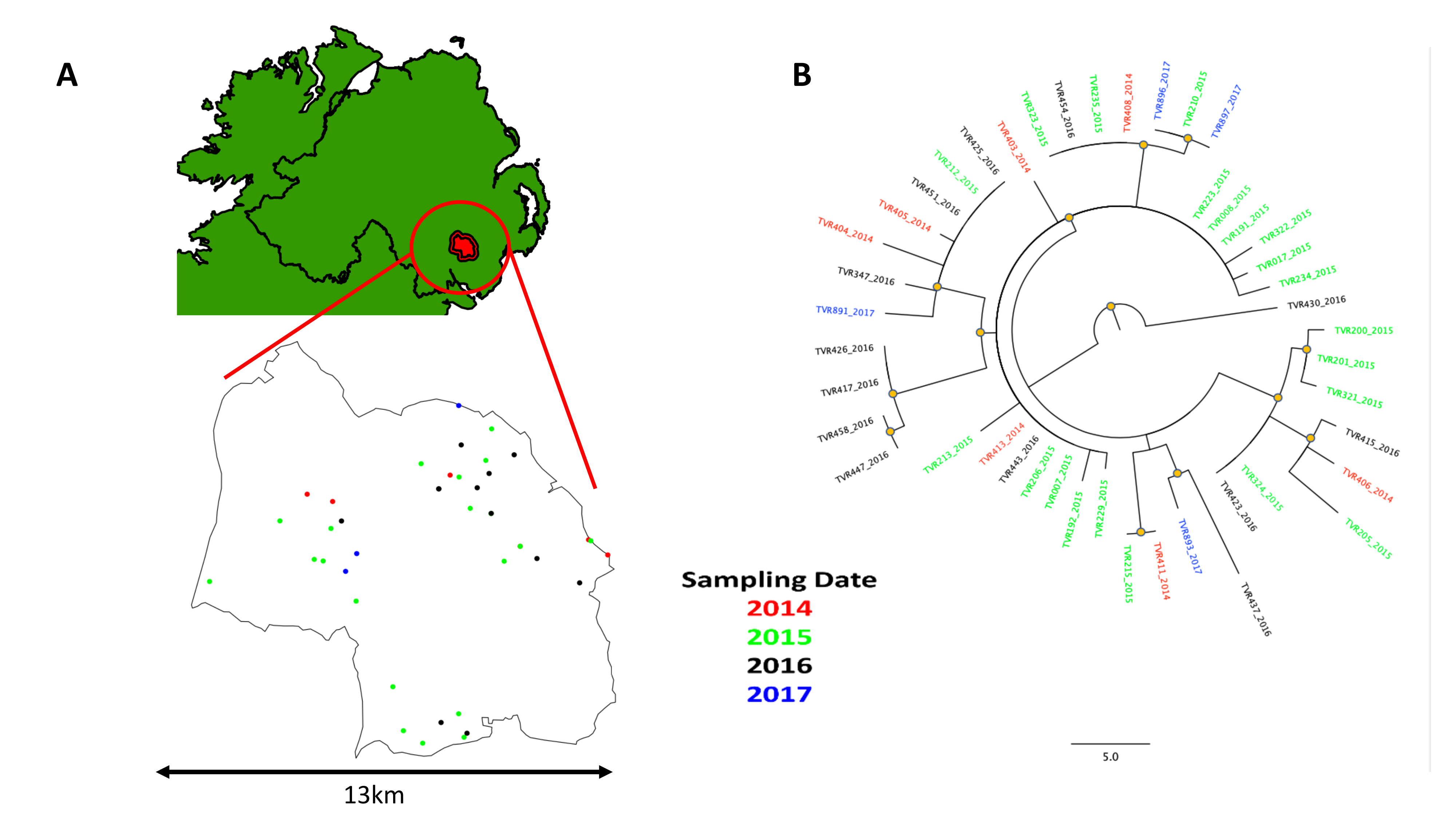

**Supplementary Figure S1** – A: Locations of 45 *M. bovis* culture positive badgers by year. B: Maximum likelihood phylogeny of 45 *M. bovis* endemic lineage isolates from TB positive badgers.

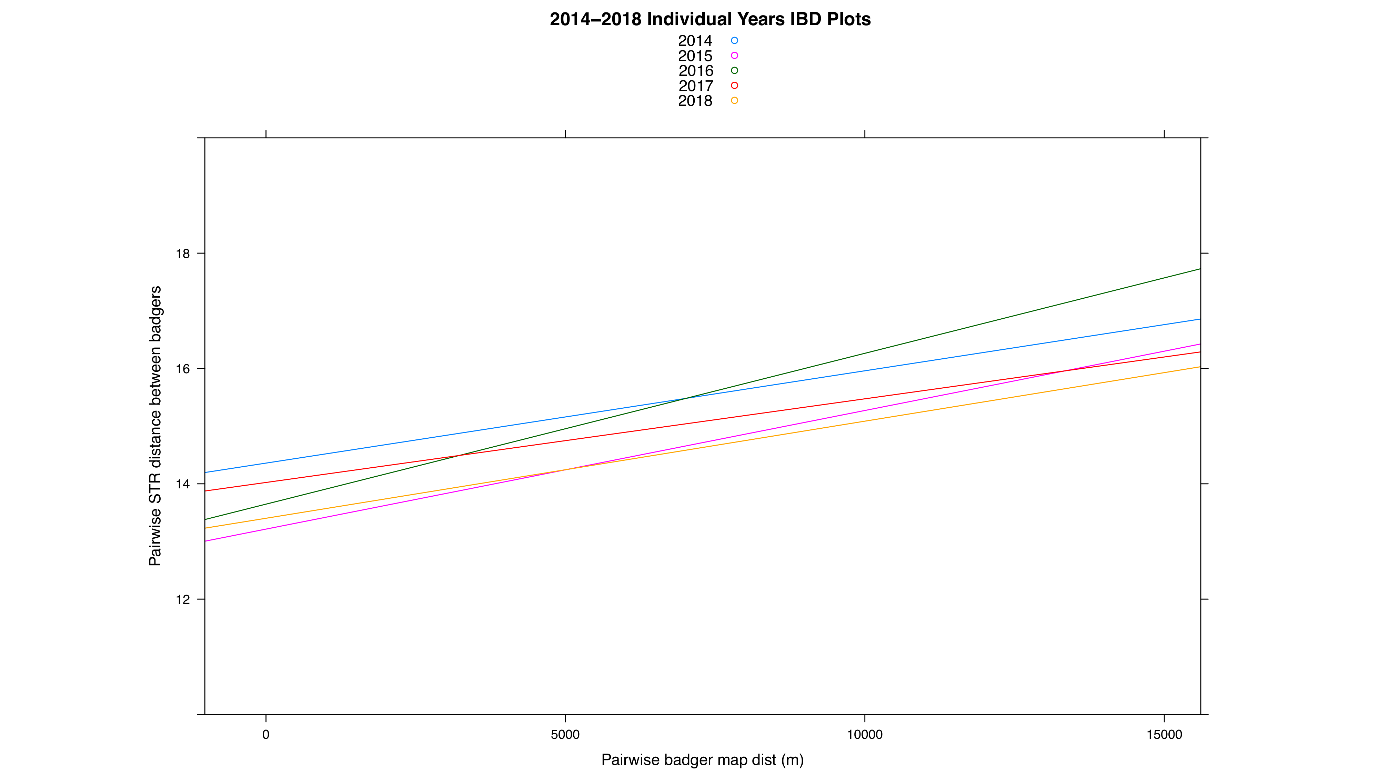

**Figure S2 -** Linear regressions of badger IBD relationships – pairwise microsatellite / STR genetic distance vs Euclidean distance for all capture years.

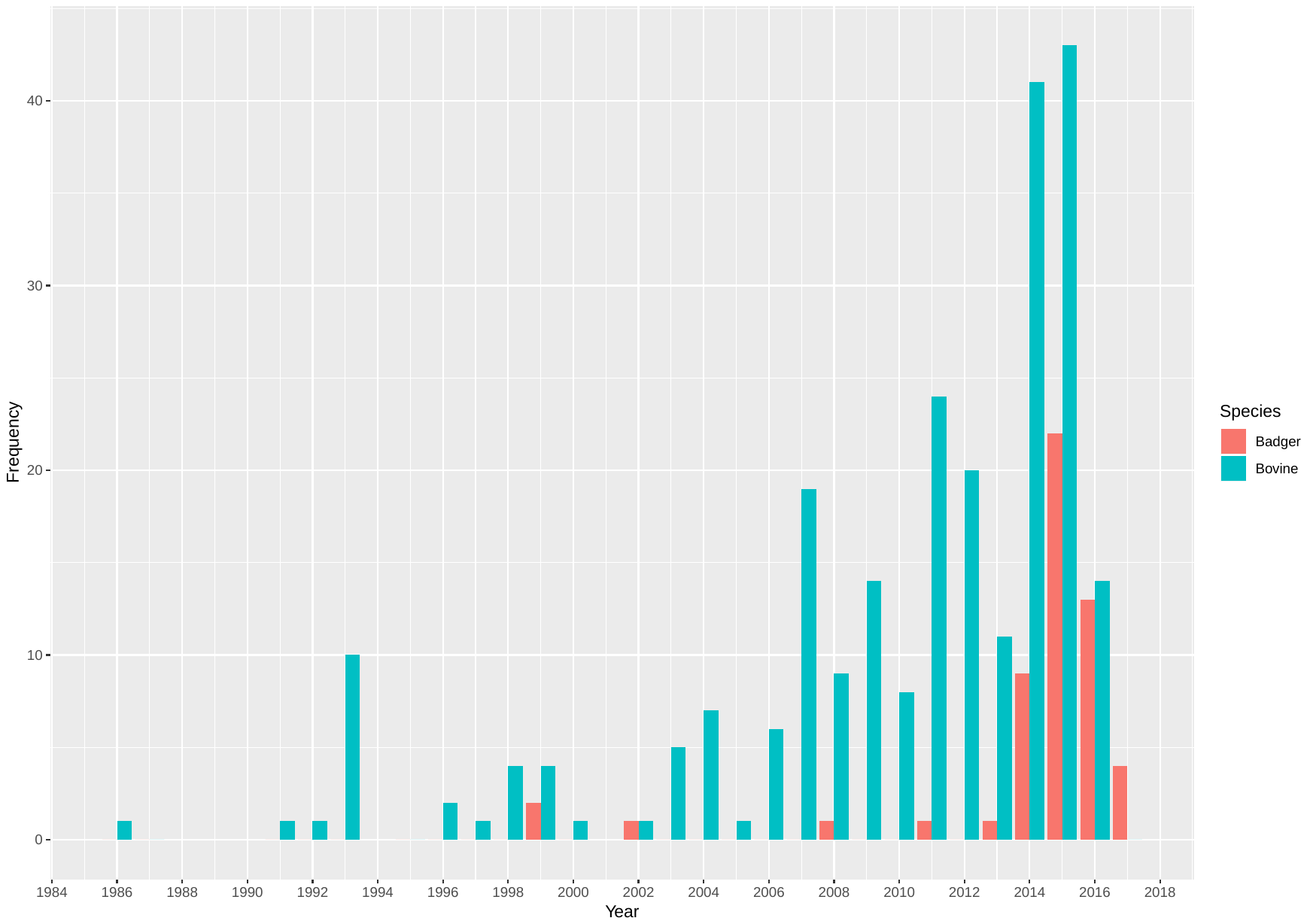

**Supplementary Figure S3 –** Sampling frequency across all years by host for endemic 6.263 lineage.

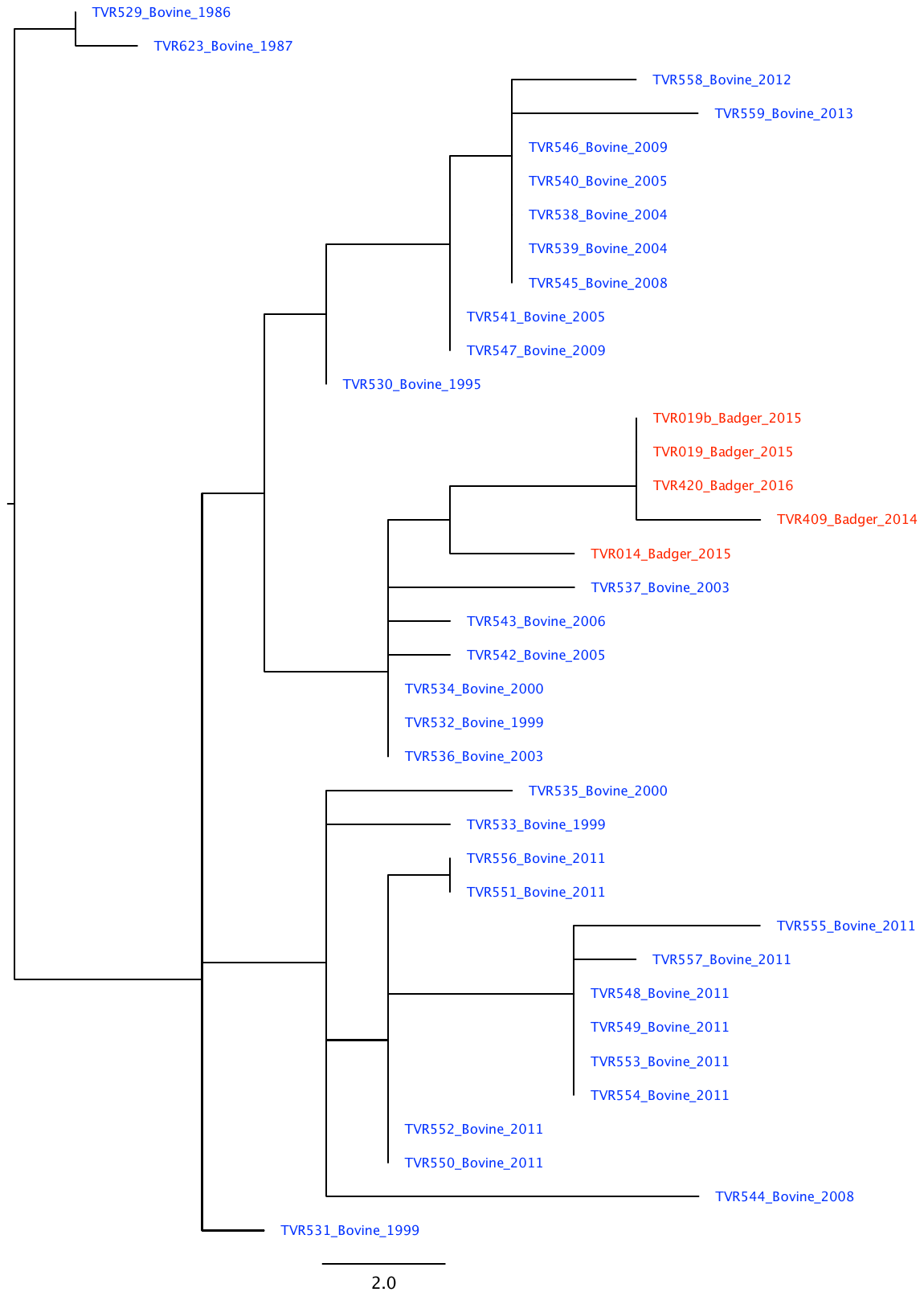

**Supplementary Figure S4 -** Maximum likelihood phylogeny (53 SNPs) of non-endemic lineage 20.131.

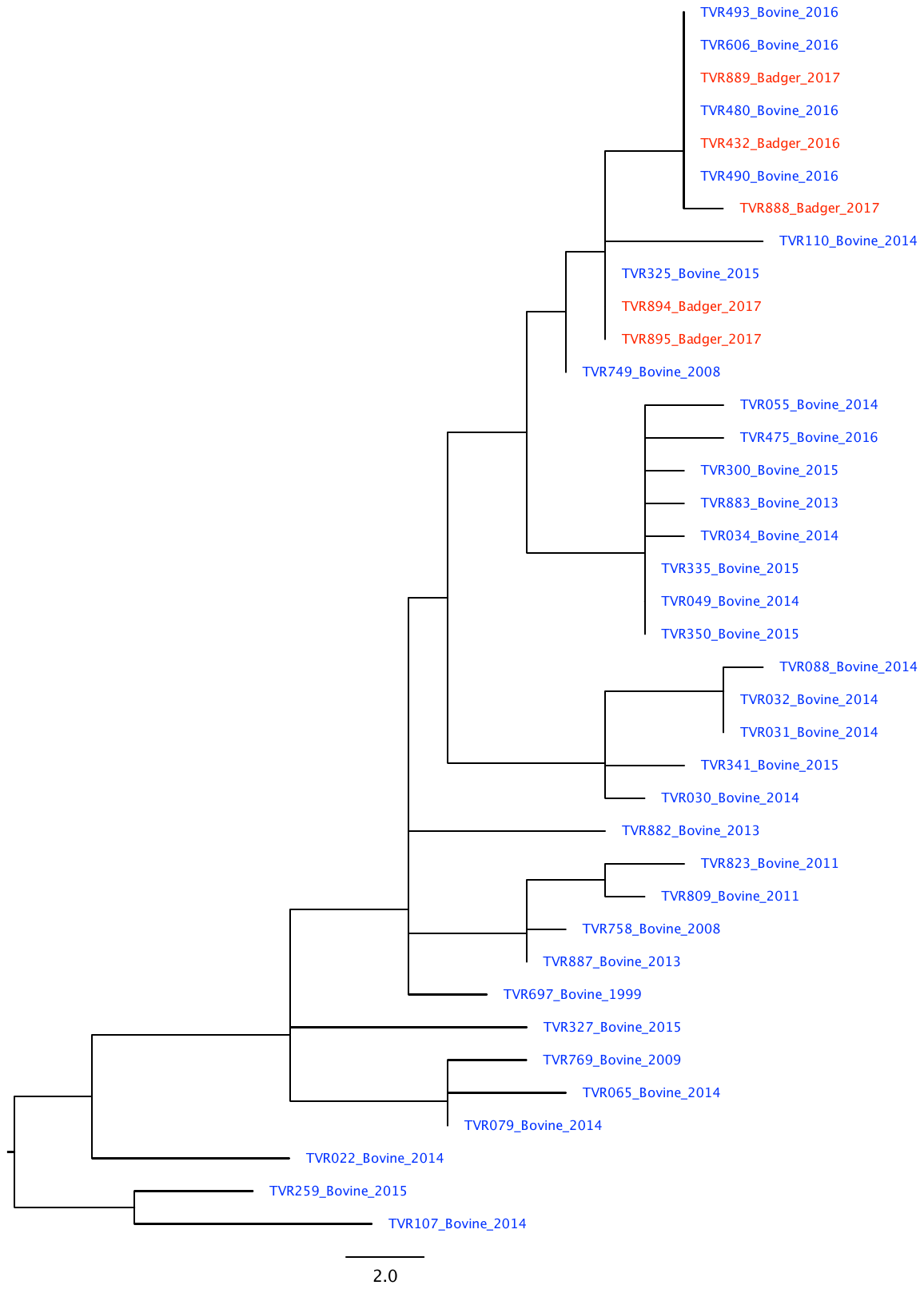

**Supplementary Figure S5 –** Maximum likelihood phylogeny (92 SNPs) of non-endemic lineage 4.140.

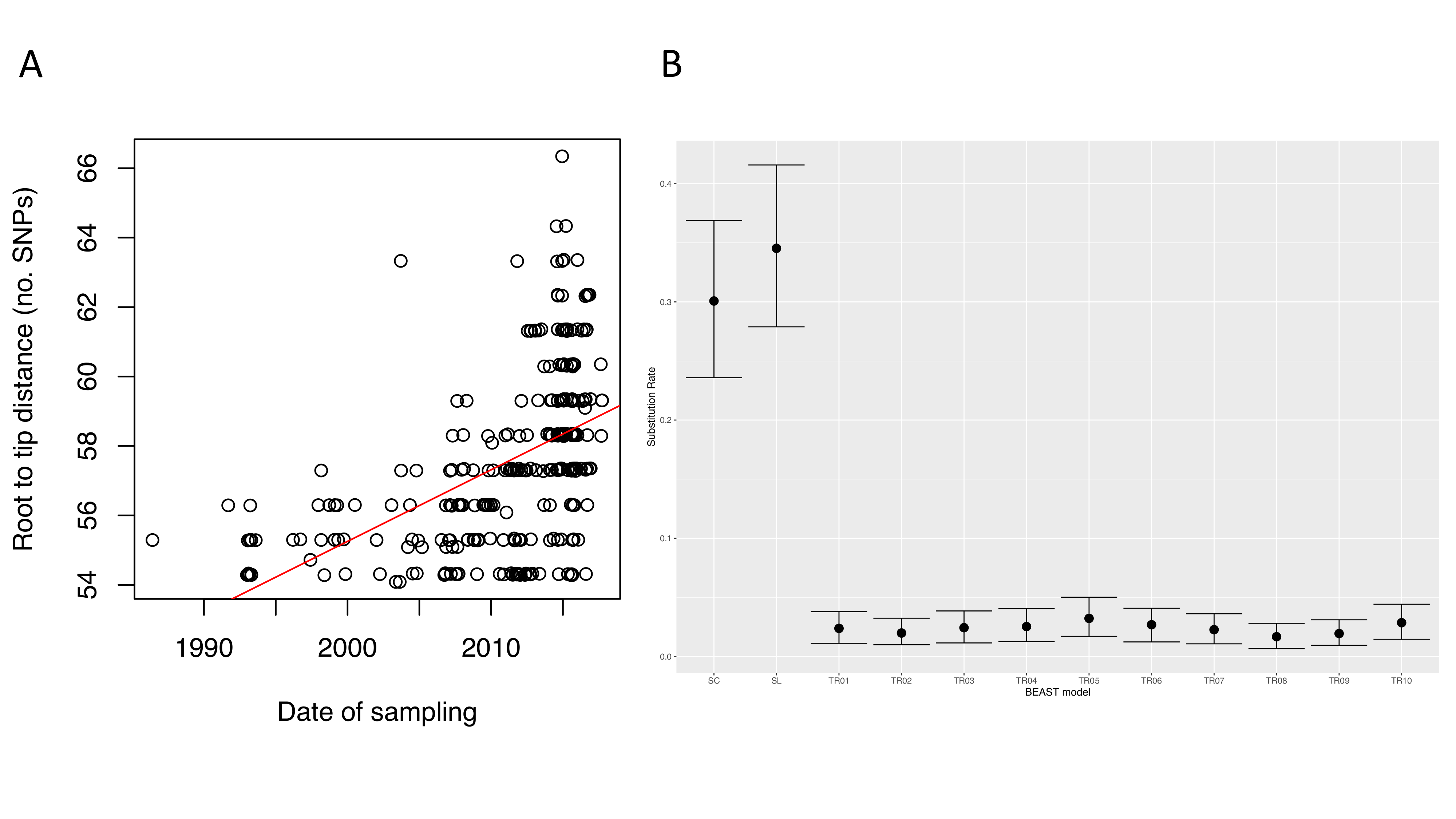
**Supplementary Figure S6 -** A: Tempest root to tip distance for n=302 isolates from endemic 6.263 lineage (R^2^ 0.25, p<0.001, beta=0.22). B: Substitution rates and 95% HPD for various models run in BEAST. SC – simple coalescent model, constant population size; SL – Skyline model; Tip date randomisation runs (TR01-TR10) with simple coalescent model.

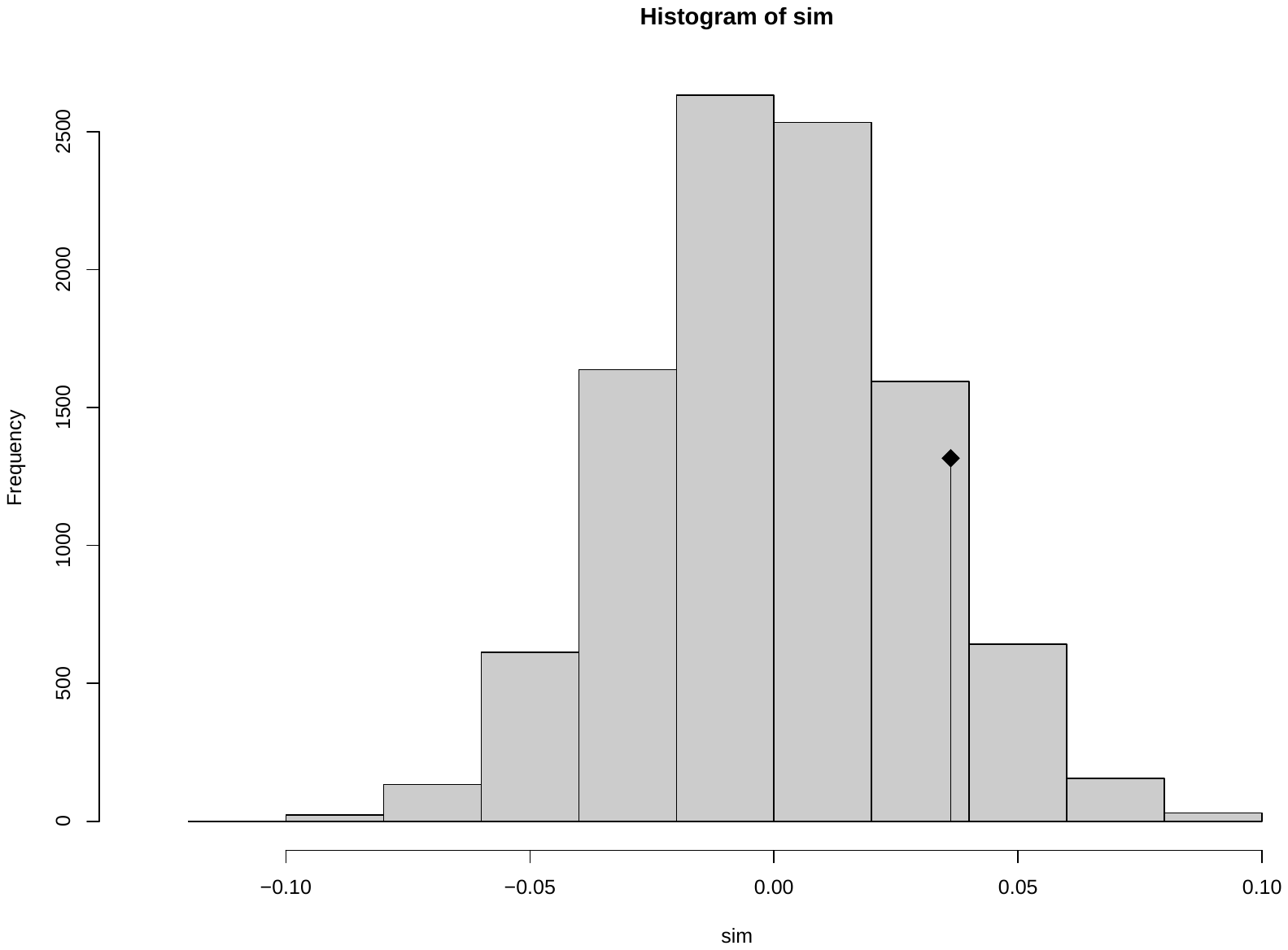

**Supplementary Figure S7 –** Mantel test of endemic clade (n=302) *M. bovis* inter-isolate SNP and Euclidean distances. r= 0.04, p=0.157

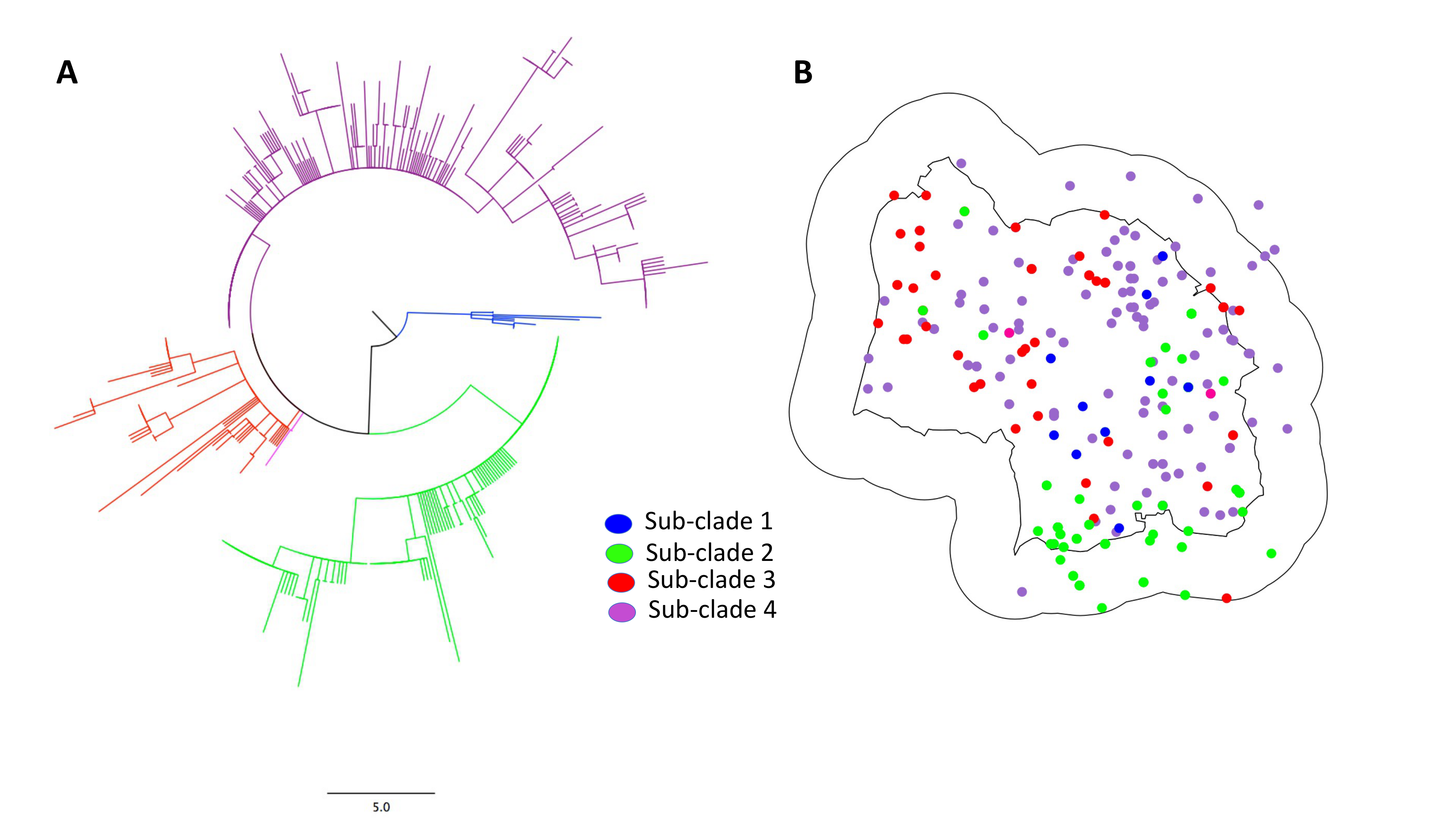

**Supplementary Figure S8** – A: maximum likelihood phylogeny of endemic clade with four major sub-clades highlighted. B: Spatial distribution of four major sub-clades across TVR zone.
