## Supplementary Materials for "Genomic epidemiology of *Mycobacterium bovis* infection in sympatric badger and cattle populations in Northern Ireland"

**Sampling representativeness of underlying bTB dynamics.**

During the 2014-2017 temporal window, the most systematic and equal effort possible, was made to detect and sample *M. bovis* from cattle and badgers. In total, 261 positive TB cultures were available from individual cattle. In badgers, we had 54 positive cultures from this period.

How representative is this sampling likely to have been? Given existing knowledge of host density in the region, known bTB prevalence in hosts and success of culture from diagnostic test positive cattle, we can try to determine an answer to this question.

**Cattle**

The TVR region is found in the Newry Divisional Veterinary Office (DVO) in County Down.

DVO area = 1466 km^2^

No. of animals in DVO = 220780

Density of animals in DVO = 150 animals per km^2^

During the TVR intensive sampling period for cattle, in which genome preps were collected from cattle (2014-2016), sampling was conducted in both the 100 km^2^ zone, and a 102 km^2^ buffer. Therefore, in total, a 202 km^2^ sampling area for cattle.

Given the density of animals noted above, the total cattle in the sampling was = 202 x 150 = 30,300.

DAERA statistics (Anon, 2019) suggest that the average animal level incidence of bTB in the Newry DVO region over the sampling period was 1.2%.

Therefore, per annum one would expect to find ~360 skin test positive animals.

Over the 2 year intensive period of cattle sampling (2014-2016), we could have expected ~720 skin test reactors.

Given a ~40% culture success rate (Robin Skuce, personal communication) from positive animals, we would have expected to get 0.4 x 720 genomes from the sampling = 288 genomes.

The latter compares favourably to the 261 genomes actually sampled during the study sampling period.

**Badgers**

Badger density in Co. Down has been observed to be ~4 animals per km^2^ (Reid et al. 2012)**.**

In the intensive period of systematic sampling (2014-2017), badgers were only taken from the 100 km^2^ study zone, not the buffer zone.

Therefore ~400 animals were available per annum for potential trapping and sampling in the study area.

Given a conservative badger bTB prevalence of ~15% (Courcier et al 2018), this suggests approximately 60 bTB positive animals per annum were potentially available for capture.

Over the three-year period of the sampling for badgers, this suggests 180 positive animals were potentially available during the study period.

Given a reported test sensitivity of 0.50 for the DPP sett side test used for badger bTB diagnosis (Courcier et al. 2020), this suggests ~90 animals.

Further, given a reported culture confirmation rate of DPP positive animals of 50.4% (Fraser Menzies, personal communication), this suggests we could have expected to get 0.504 x 90 genomes from the sampling = 45 genomes.

This also compares favourably to the 54 positive cultures actually sampled in the study period.

**Conclusions.**

The variable sensitivities of the bTB diagnostic tests employed in both cattle (de la Rua Domenech et al. 2006) and badgers (Chambers et al 2009; Courcier et al. 2020) mean that a significant proportion of truly infected animals are missed. Additionally, those that are detected will not all yield culturable *M. bovis* isolates. On the wildlife side, we know that the phenomenon of badger ‘trappability’ means that some animals are more likely to be captured than others (Tuyttens et al. 2001; Byrne et al. 2012). These factors mean that there is a proportion of the epidemic that is unobserved. Indeed, our estimates of interspecies pathogen transitions explicitly incorporate the latter fact by being conservative in nature, an acknowledgement that in the coalescing lineages we observe, we are likely missing undetected hosts and therefore underestimating transition rates.

However, the longitudinal, systematic structure of the sampling undertaken here should minimise the risk of making spurious inferences. Cattle bTB data collected in Northern Ireland result from annual testing of all animals, unlike some regions in Britain that employ protocols of differing frequencies of testing depending on prevalence (Allen et al. 2018). The wildlife trapping and sampling undertaken here is denser and more contemporaneous with cattle sampling. Such wildlife sampling is potentially more representative of the badger population than has previously been achieved by using animals killed in road traffic accidents (RTAs) (Biek et al. 2012; Trewby et al. 2016). Exclusive use of RTA animal isolates may introduce bias to datasets on account of them primarily being taken from animals undertaking extra-territorial dispersion for mating purposes (Roper, 2010). Inter-annual mark recapture analyses in the TVR zone indicate a mean annual recapture rate of 56.2%, which compares favourably to similar exercises undertaken elsewhere in Ireland (Byrne et al. 2012).

The rough projections above suggest that numbers of expected samples collected from both hosts, given prior knowledge of host density in landscape, bTB prevalence, test sensitivity and culture confirmation rates, are largely in line with actual observed sample numbers. Given the prior information available, one would expect to collect more cattle samples than badger samples with the difference being in the order of x4-x6.

We conclude therefore that the TVR sampling is likely to be representative of the weight of infection and diversity of pathogen in both hosts.
