## Supplementary Tables for "Genomic epidemiology of *Mycobacterium bovis* infection in sympatric badger and cattle populations in Northern Ireland"

| **Year** | **N** | **Mean Na** | **Mean He** | **Mean Ho** | **Mean Fis** | **Mean AR** |
| --- | --- | --- | --- | --- | --- | --- |
| **2014** | 273 | 4.9 | 0.52 | 0.49 | 0.06 | 4.77 |
| **2015** | 152 | 5.0 | 0.51 | 0.49 | 0.04 | 4.81 |
| **2016** | 97 | 4.6 | 0.52 | 0.50 | 0.04 | 4.53 |
| **2017** | 113 | 5.1 | 0.52 | 0.48 | 0.09 | 5.05 |
| **2018** | 134 | 4.8 | 0.51 | 0.49 | 0.04 | 4.73 |
| **ALL YEARS** | 769 | 4.9 | 0.52 | 0.49 | 0.05 | 4.78 |

**Table S1** – Badger meta population genetic summary statistics averaged across 14 microsatellite loci. N = no. of animals genotyped successfully; Na = no. of alleles observed per locus; He = expected heterozygosity; Ho = observed heterozygosity; Fis = fixation index (level of inbreeding per locus); AR = Allelic richness.

| **Model** | **Variable** | **Slope** | **P value** |
| --- | --- | --- | --- |
| **SNP dist ~ Euclidean dist + time of isolation difference + microsat dist** | Euclidean distance | 2.83x10^-4^ | 0.04* |
|  | Time difference | 8.81x10^-1^ | 0.08 |
|  | Microsatellite distance | 2.92x10^-2^ | 0.76 |
| **Full model R^2^ = 0.04**  **P value = 0.10**  **F = 14.16** | | | |

**Table S2 -**  Multiple regression on distance matrices (MRM) analysis of *M.bovis* inter-isolates SNP distance vs pairwise Euclidean distance between +ve badgers, pairwise time difference between *M. bovis* isolations and pairwise microsatellite genetic distance between host badgers. * = p<0.05.

|  | Biek et al 2012 | Trewby et al 2016b | Crispell et al 2017 | Salvador et al 2018 | Crispell et al 2019 |
| --- | --- | --- | --- | --- | --- |
| Substitution rate | 0.15 | 0.20 | 0.53 | 0.37 | 0.28 |

**Table S3** – *M. bovis* substituton rates (substitutions per genome per year) as assessed in previous studies.
